# Sexual dimorphism in epigenomic responses of stem cells to extreme fetal growth

**DOI:** 10.1101/008482

**Authors:** Fabien Delahaye, N. Ari Wijetunga, Hye J. Heo, Jessica N. Tozour, Yong Mei Zhao, John M. Greally, Francine H. Einstein

**Affiliations:** 1300 Morris Park Avenue, Block Building, Room 631, Department of Obstetrics & Gynecology and Women’s Health, Albert Einstein College of Medicine, Bronx, NY, United States, 10461; 1301 Morris Park Avenue, Price Building, Room 322, Department of Genetics, Albert Einstein College of Medicine, Bronx, NY, United States, 10461

## Abstract

Extreme fetal growth is associated with increased susceptibility to a range of adult diseases through an unknown mechanism of cellular memory. We tested whether heritable epigenetic processes in long-lived CD34+ hematopoietic stem/progenitor cells (HSPCs) showed evidence for re-programming associated with the extremes of fetal growth. Here we show that both fetal growth restriction and over-growth are associated with global shifts towards DNA hypermethylation, targeting *cis*-regulatory elements in proximity to genes involved in glucose homeostasis and stem cell function. A sexually dimorphic response was found, intrauterine growth restriction (IUGR) associated with substantially greater epigenetic dysregulation in males but large for gestational age (LGA) growth affecting females predominantly. The findings are consistent with extreme fetal growth interacting with variable fetal susceptibility to influence cellular aging and metabolic characteristics through epigenetic mechanisms, potentially generating biomarkers that could identify infants at higher risk for chronic disease later in life.

## INTRODUCTION

Environmental factors have the potential for significant impact on normal development and health throughout the life span. Suboptimal intrauterine conditions represent a specific type of environmental exposure that is associated with increased risk for cardiovascular disease^1,2^ and premature death in adulthood^3^. A substantial amount of evidence has also demonstrated the relationship between poor maternal nutrition or low birth weight with a range of metabolic disorders and obesity in humans^4,5^ with animal studies further corroborating these findings^6^. At the opposite end of the spectrum of extreme fetal growth, excess nutrition leading to large for gestational age (LGA) birth weights is associated with similar adult phenotypes, with increased risk for premature mortality^3^ and a range of other age-related diseases^7^. Fetal growth restriction and over-growth show a decline in resistance to chronic disease in adulthood and involvement of multiple organ systems, which is typical of normal aging and may represent a precocious aging phenotype associated with both extremes of the fetal growth spectrum^8^.

Adverse exposures appear to be particularly consequential in early life, possibly due to the rapid expansion of cell populations necessary for growth, and the dynamism of cellular differentiation and lineage commitment that occurs during this period of development. Inherent to the differentiation process is the modification of transcriptional regulatory patterns. These include epigenetic regulators that are capable of transmitting newly established regulatory marks through cell replication^9^. Environmentally induced perturbations of the cell’s normal epigenetic regulatory controls may be maintained in long-lived, self-renewing cells, maintained through proliferation and resulting in functional consequences later in life. While alterations in DNA methylation has been associated with the cumulative exposures inherent to aging^10^, environmental exposures early in life may induce addition dysregulation of the epigenome conferring increased susceptibility for age-related disease at a younger age.

We^11,12^ and others^13–15^ have explored the possibility that non-random epigenetic changes are associated with IUGR. In studies of disease or phenotype-associated epigenetic changes, the choice of cell type generally represents a compromise between accessibility, purity, quantity and mechanistic relevance. Unpurified peripheral blood leukocytes have previously been studied in individuals whose mothers were exposed prenatally to famine. Altered DNA methylation of multiple sites within the differentially methylated region (DMR) of the imprinted *Insulin-like growth factor 2* (*IGF2)* gene was found in subjects decades later^13^. Cord blood leukocytes have also been used to demonstrate associations of DNA methylation with *in utero* conditions and birth weight^15,16^. Another commonly studied tissue type is the placenta, which functions at the maternal-fetal interface and may be a potential mediator of intrauterine environmental conditions^17–19^. However, testing the placenta does not address the latent risk in adulthood of chronic disease, which has to be mediated by somatic cells of the offspring. Furthermore, the use of samples of mixed populations of cells in DNA methylation studies, such as those sampled from highly heterogeneous placental tissue, is now recognized as a major source of experimental artifact that limits interpretability of results^20,21^.

We focus on hematopoietic stem/progenitor cells (HSPCs), purified using the CD34 surface marker to reduce cell subtype effects^20^. HSPCs include a subset of long-term, self-renewing stem cells that persist through the life of the individual^22^, allowing the cellular propagation or the ‘memory’ of exposure to temporally remote suboptimal conditions. The role of CD34+ HSPCs in the maintenance of vascular integrity^23,24^ is mechanistically relevant for the adult phenotype associated with increased risk for cardiovascular disease^4,25^. We studied infants born with the two extremes of fetal growth, IUGR and LGA, compared with control infants with appropriate weight for gestational age. Due to the thorough characterization of CD34+ HSPCs by the Roadmap Epigenomics Program, we were able to exploit the mapping of chromatin constituents to define empirically the *cis*-regulatory elements, such as promotors and enhancers, specific to this cell type (Wijetunga, Delahaye *et al.*, manuscript co-submitted), allowing us to interpret changes in DNA methylation at otherwise unannotated loci in the genome.

## RESULTS

### Genome-wide DNA methylation profiling

We perform genome-wide DNA methylation profiling on purified CD34+ HSPCs from 60 subjects, 20 in each of three groups defined by appropriate or excessively large or small birth weight and ponderal index for gestational age and sex (Table 1). The HELP-tagging assay is used as a survey technique testing ~1.8 million loci quantitatively at nucleotide resolution and including relatively CG dinucleotide depleted loci^26^. This assay generates a methylation score that is inversely correlated to DNA methylation level, with a methylation score of 0 indicating full methylation and 100 indicating complete lack of methylation, based on a normalized ratio between tag counts generated by the methylation-sensitive enzyme HpaII and its methylation-insensitive isoschizomer MspI^27^. Based on quality control measures (**Methods** and **Supplementary Fig. 1**), 993,514 loci are selected for further analyses. Of these, 10,043 loci are defined as candidate differentially-methylated loci using batch-adjusted significance and degree of methylation difference thresholds in comparisons of IUGR and LGA infants with the normal birth weight controls. We observe a global relative shift towards DNA hypermethylation in CD34+ HSPCs in both IUGR and LGA subjects when compared with the controls (Fig. 1a). The clustering of cases (LGA/IUGR) is not uniform, with a subset of cases clustering with controls (Fig. 1b), indicating that epigenetic dysregulation does not occur universally as a response to extreme fetal growth. While there exists a subset of common loci altered in both IUGR and LGA neonates, most of the dysregulated loci are distinctive between these groups (Fig. 1c,d). We also see an overlap of <u>genes</u> (as opposed to loci) undergoing differential DNA methylation (**Supplementary Fig. 2**).

**Figure 1.**
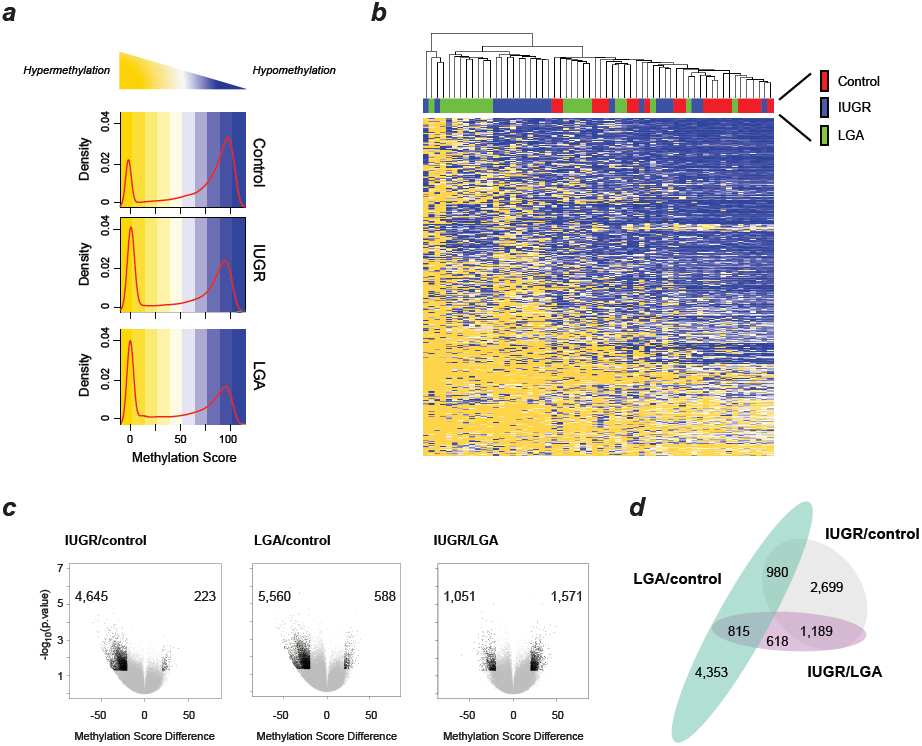
Genome-wide DNA methylation profiles (**a**) Density plots of methylation scores for IUGR or LGA compared with controls. The distributions of DNA methylation scores are shown in red. (**b**) A self-organizing heatmap of candidate differentially methylated loci showing clustering by sample. (**c**) Volcano plots of DNA methylation score differences for IUGR compared with control, LGA compared with control and IUGR compared with LGA, based on 993,514 loci throughout the genome. Differentially methylated loci with p value <0.05 and methylation difference >|20| are shown in black. (**d**) Differentially methylated loci meeting threshold criteria are quantified in a proportional Venn diagram for each comparison.

**TABLE 1.** Clinical cohort characteristics.

|  | COHORT 1 |  |  | COHORT 2 |  |  |
| --- | --- | --- | --- | --- | --- | --- |
|  | Control<br>(n=20) | IUGR<br>(n=20) | LGA<br>(n=20) | Control<br>(n=8) | IUGR<br>(n=8) | LGA<br>(n=8) |
| **Gestational age, weeks** | 39.3 ± 0.3 | 38.7 ± 0.4 | 39.2 ± 0.2 | 39.1 ± 0.5 | 38.7 ± 0.4 | 39.8 ± 0.3 |
| **Birthweight, g** | 3,150 ± 64 | 2,515 ± 69 <sup>a</sup> | 3,996 ± 81 <sup>a</sup> | 3,150 ± 161 | 2,498 ± 78 <sup>b</sup> | 4,053 ± 67 <sup>a</sup> |
| **Ponderal index, g/cm<sup>3</sup>** | 2.8 ± 0.03 | 2.3 ± 0.04 <sup>a</sup> | 3.2 ± 0.03 <sup>a</sup> | 2.8 ± 0.07 | 2.3 ± 0.04 <sup>a</sup> | 3.4 ± 0.07 <sup>a</sup> |
| **% Male** | 50 | 50 | 50 | 50 | 50 | 50 |
| **Maternal age, years** | 24.6 ± 0.9 | 26.5 ± 1.3 | 31.1 ± 1.1 <sup>a</sup> | 33.1 ± 1.7 | 27.7 ± 2.6 <sup>b</sup> | 31.5 ± 1.6 |
| **Pre-pregnancy BMI,**<br>**kg/m<sup>2</sup>** | 26.6 ± 1.3 | 25.5 ± 0.9 | 29.0 ± 1.3 | 27.0 ± 2.1 | 26.5 ± 2.8 | 28.5 ± 1.4 |
| **Weight gain, pounds** | 29.4 ± 2.7 | 23.0 ± 2.7 | 29.9 ± 2.1 | 27.0 ± 5.1 | 12.7 ± 5.1 | 24.8 ± 2.9 |
|  |  |  |  | a = p<0.001 compared with<br>Control |  |  |
|  |  |  |  | b = p<0.05 compared with<br>Control |  |  |

Showing mean ± standard deviation values.

### Sexual dimorphism associated with the extremes of fetal growth

Sex-specific comparisons for DNA methylation patterns are shown between control and IUGR and LGA subjects (Fig. 2). Both IUGR males and females show a shift in DNA methylation profiles compared to controls, but the number of hypermethylated loci is markedly higher in males compared to females (Fig. 2a). Sex-specific differences are also seen in the comparison of LGA to controls, with LGA females showing an increase in the overall number of candidate differentially-methylated loci compared to males (Fig. 2b). These findings indicate a sexual dimorphism in the epigenetic responses of HSPCs to the extremes of growth conditions *in utero*.

**Figure 2.**
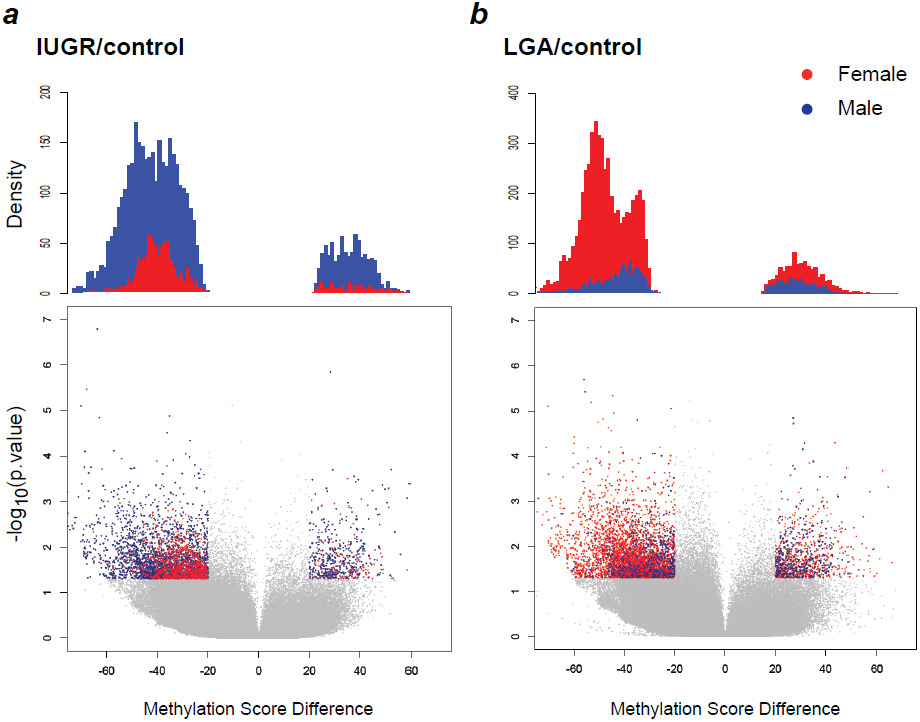
Sexual dimorphism in IUGR males and LGA females for differentially methylated loci The lower panels show volcano plots of DNA methylation score differences, the upper panels quantify the densities of differentially methylated loci (p value<0.05 using ANOVA with pairwise two-tailed Tukey-tests, methylation difference >|20|). (**a**) IUGR compared with controls, (**b**) LGA compared with controls.

### Targeting of DNA methylation changes to specific genomic contexts

While the consequences of DNA methylation changes at recognized promoter sequences are generally predictable, a genome-wide study of this type can generate a majority of findings in un-annotated genomic locations. To predict the functional consequences of these candidate differentially-methylated loci, we take advantage of the mapping of chromatin components in CD34+ HSPCs performed as part of the Roadmap Epigenomics Program. The details of this annotation are described in a separate report (Wijetunga, Delahaye *et al.*, manuscript in review) and involve the use of the Segway algorithm^28^ to generate genomic features (**Methods**) that are then interpreted using Self-Organizing Maps^29^. We are thus able to define candidate promoters, enhancers, transcribed sequences and repressive chromatin in the epigenome specific to the CD34+ HSPC population. Every HpaII site is then assigned to a candidate feature based on its genomic position. The HELP-tagging assay represents each of the candidate genomic features (based on 993,514 loci) and the candidate differentially-methylated loci (10,043) are significantly enriched in Segway features 4 (enhancers, p<0.001) and 6 (promoters, p<0.001), indicating preferential targeting to transcriptional regulatory elements (Fig. 3a). We show an example of the mapping of the one the candidate differentially-methylated loci, to the promoter of the *Retinoid X receptor, alpha* (*RXRA)* gene, at an annotated CpG island, and within the Segway feature 6 annotation indicating candidate promoter function. The HELP-tagging derived methylation scores for cases (IUGR and LGA combined) are compared with controls to demonstrate the magnitude of the change at this locus (Fig. 3b).

**Figure 3.**
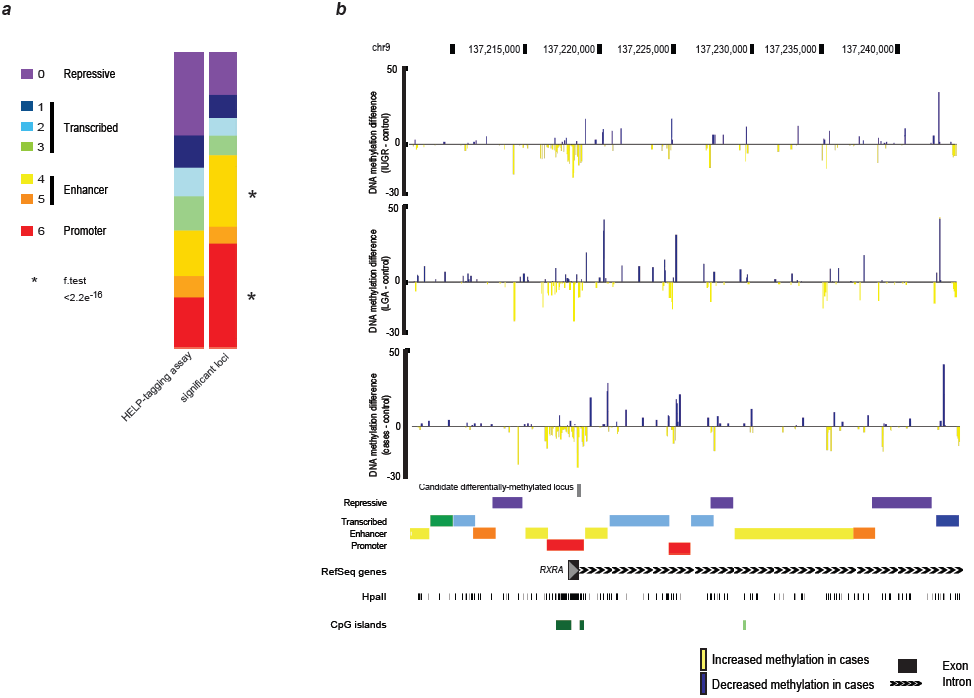
Candidate differentially-methylated loci are enriched at *cis*-regulatory elements (**a**) Based on empirical annotation of promoter, enhancer, repressive and transcribed regions, enrichment of candidate differentially-methylated loci (n=10,043) in cases (IUGR and LGA) compared with controls is illustrated with significance values shown for enriched sequence features. The bar on the left represents the proportional representation of each feature in terms of loci tested by HELP-tagging, while the bar on the right shows the proportions of features at which differentially-methylated loci are found. Significant enrichment for differential methylation at candidate promoters and enhancers is observed. (**b**) An example of the *RXRA* gene with a candidate differentially-methylated locus is shown. The DNA methylation score differences between controls and IUGR (top), LGA (middle) and cases (bottom, IUGR and LGA combined) is depicted, with a site identified as being a candidate differentially-methylated locus in the CpG island promoter region shown in gray. Blue, positive values represent decreased DNA methylation in the cases of extreme fetal growth, yellow, negative value increased methylation.

### Targeting of DNA methylation changes to genes with specific properties

We test whether the subset of loci affected by DNA methylation changes are enriched at a specific subset of genes characterized by concordance of function of their protein products. A candidate differentially methylated locus is linked to a specific gene if the site is (a) located in proximity to the transcription start site of the RefSeq gene and (b) overlapping candidate regulatory loci (features 4 or 6). We select only those candidate promoters (feature 6) within ±2 kb and candidate enhancers (feature 4) within ±5 kb of RefSeq transcription start sites. While enhancers can act over substantially longer distances than 5 kb^30^, we are deliberately conservative in restricting the distance so that we would be more likely to associate an enhancer with the gene upon which it exerts its effects. The resulting list of genes is used to perform a gene set enrichment analysis (GSEA). Traditional GSEA does not take into account the physical characteristics of the gene and has been shown to be biased by factors such as the numbers of CG dinucleotide sites associated with different classes of gene and gene promoters^31^. To address this, the Bioconductor package *GoSeq*^32^ was developed to control for variability of length of genes. We adapted *GoSeq* to normalize our data to control for the number of CG dinucleotides linked to each gene by the above criteria. Detailed information describing the results of the normalized GSEA is shown in Supplementary **Tables 1-2**. Among the different significant pathways from KEGG (Kyoto Encyclopedia of Genes and Genomes), two pathways of interest emerge as significant regardless of group comparison: the KEGG pathways for *Maturity onset diabetes of the young*, relevant to glucose homeostasis and *Hedgehog (HH) signaling*. Both of these pathways contain genes involved in proliferation, differentiation and self-renewal capabilities of stem cells. Permutation analysis was performed to confirm the significance of these results. Based on the criteria for assigning HpaII sites to RefSeq genes described above, the HELP-tagging assay represents 97.6% of RefSeq genes, so we randomly select from within this group of genes the same number of genes used to define our pathways, and perform the GSEA analysis 1,000 and 3,000 times to test how frequently the same pathways are identified as, defining the significance of our detection of these pathways as p<10^−3^. The same pathways are targeted by IUGR and LGA even when the loci involved are not identical (Fig. 4). A similar effect is seen for the loci affected differentially between males and females (Supplementary **Fig. 3**). These findings combine to show convergence of dysregulation of the same pathways by IUGR and LGA and in male and female subjects respectively, even though the loci targeted for DNA methylation changes are not necessarily the same in each group.

**Figure 4.**
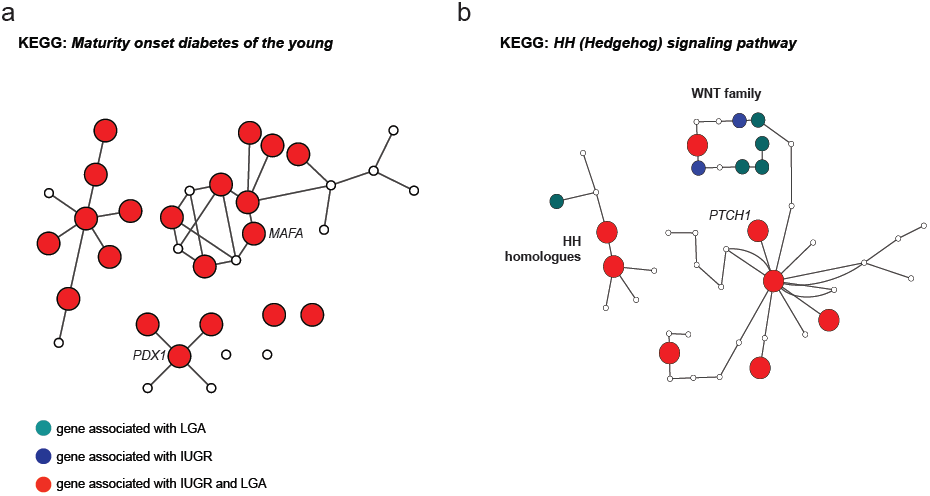
Network Analysis A network representation of KEGG pathways for (**a**) *Maturity onset diabetes of the young* and (**b**) *Hedgehog (HH) signaling*. Nodes are color- and size-coded based on the association of genes represented by each node with LGA or IUGR, or with both LGA and IUGR. Edges (solid lines) represent known physical interaction between genes.

### Verification and validation

To test the robustness of our genome-wide technique, we assess the reproducibility of DNA methylation differences at our candidate differentially-methylated loci using single locus quantitative validation studies. We first perform verification studies on samples from Cohort 1, on whom the genome-wide studies had been performed, testing 4 loci selected for differing levels of DNA methylation in 24 subjects. A strong correlation between bisulphite MassArray and HELP-tagging is found (R^2^ = 0.98, Supplementary **Fig. 4**). In a second, independent set of CD34+ HSPC samples (Cohort 2) consisting of 8 new subjects/group (control, IUGR, LGA) with equal numbers of males and females in each group, we perform a targeted bisulphite sequencing (TBS) assay, using bisulphite treatment, targeted PCR and massively-parallel sequencing to measure DNA methylation at 72 loci in the 24 subjects (see **Methods** and **Supplementary Table 3**). The correlations between HELP-tagging with MassArray in Cohort 1 (R^2^=0.98, **Supplementary Fig. 4**) and with TBS in Cohort 2 (R^2^=0.72, **Supplementary Fig. 5** and **Supplementary Table 3**) are both strong. These highly quantitative verification and validation studies demonstrate the technical robustness of the genome-wide HELP-tagging assay, as well as the potential to validate DNA methylation differences, even when using a new cohort of subjects. Of the 54 candidate differentially-methylated loci from the HELP-tagging group comparisons, we focus on loci implicated by our *GOseq*-normalized GSEA results, using primers for candidate differentially-methylated loci proximal to *WNT6* (Fig. 5a) and *PTCH1* (Fig. 5b) from the *Hedgehog (HH) signaling* pathway and *MAFA* from the *Maturity onset diabetes of the young* pathway (**Supplementary Fig. 6**). We find the direction of DNA methylation changes to be concordant between genome-wide and targeted assays for all three loci, with statistically significant differences demonstrable for TBS data from *WNT6* (p=0.023) and *PTCH1* (p=0.014) (**Supplementary Table 4**). We also show the *PTCH1* and *WNT6* genes to have increased DNA methylation by TBS at local *cis*-regulatory elements in cases (IUGR and LGA) compared to controls (**Supplementary Table 4**, p<0.05). Finally, we interrogate loci associated with genes that are differentially methylated on average between cases (IUGR plus LGA) and controls in Cohort 1 and previously found to have epigenetic alterations related to metabolic syndrome and type 2 diabetes mellitus, *IGF2*^33,34^ and *RXRA*^35^. A positive correlation is seen between the HELP-tagging and TBS DNA methylation levels, but the TBS DNA methylation differences between cases and controls are of insufficient magnitude for statistical significance to be attributed (**Supplementary Table 4**). The number of new samples in Cohort 2 is limited and thus these validation studies are likely to confirm loci of major effect only. Overall, the TBS data are concordant with the genome-wide data, indicating that the conclusions based on the genome-wide results are tenable.

**Figure 5.**
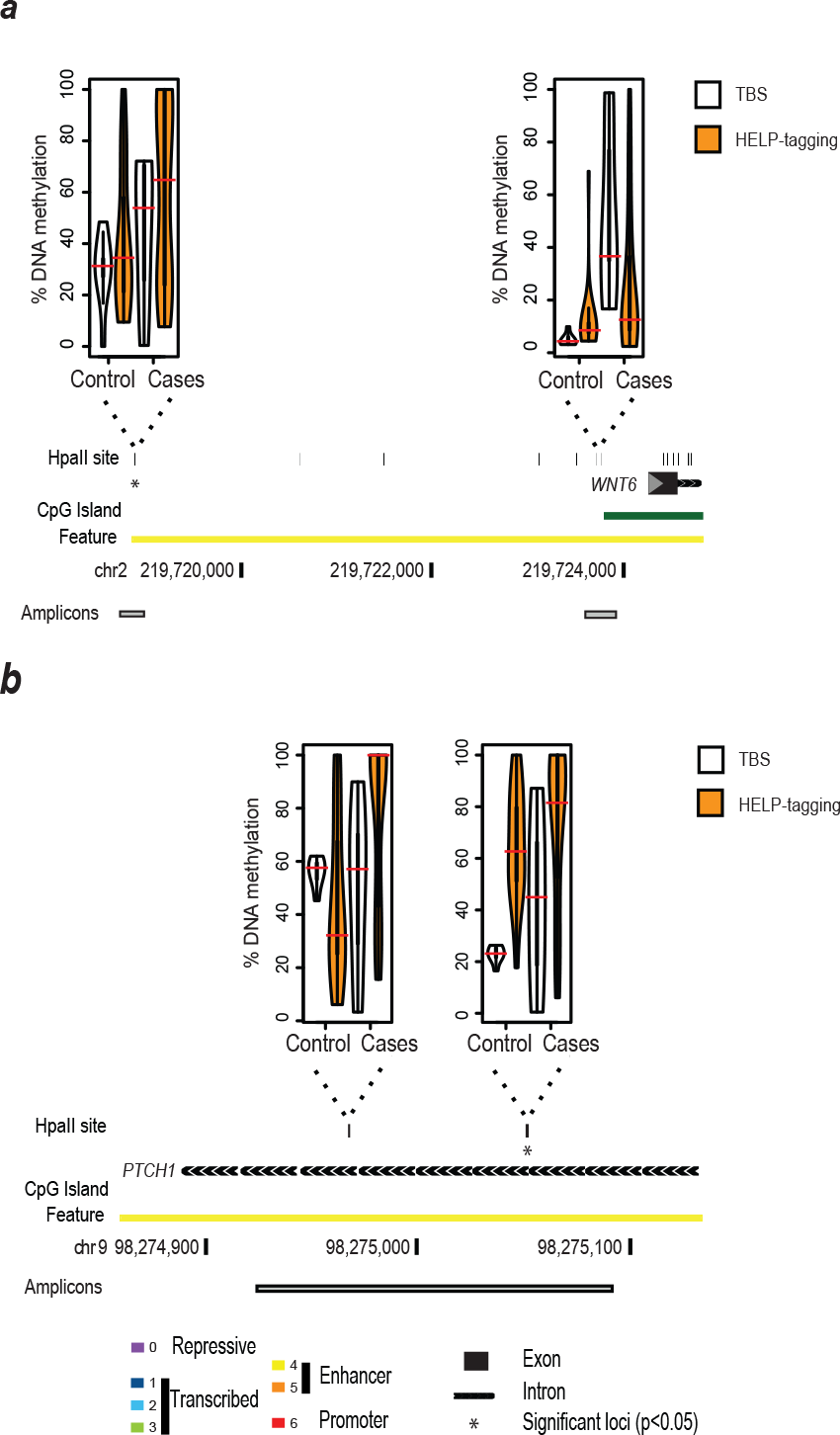
Biological validation Validation of significant loci of interest by targeted bisulphite sequencing (TBS) in Cohort 2 for loci at the (**a**) *WNT6* and (**b**) *PTCH1* genes. Candidate differentially-methylated loci are shown as the HpaII sites within the amplicon regions (gray boxes), with results of DNA methylation distributions for controls and cases (IUGR and LGA combined) from HELP-tagging (orange) and TBS (white) depicted as violin plots (mean shown in red, 1^st^ and 3^rd^ quartile are depicted by the thick black bar). The results show concordance for similar types of changes between HELP-tagging and TBS results at these loci with significant results (p<0.05, t-test) for TBS marked with asterisks.

## DISCUSSION

Here we show for the first time epigenetic changes associated with the two very different types of intrauterine conditions reflected by extremes of fetal growth. All subjects were healthy, full-term neonates without any anomalies or dysmorphic features that would suggest an etiology innate to the fetus. We combined birth weight with ponderal index values to define the distinct groups of infants at either end of the fetal growth continuum. Both groups with abnormal growth demonstrate global shifts of increased DNA methylation compared to appropriately grown neonates. Although the underlying differences in environmental exposures cannot be measured with precision in these subjects, the intrauterine conditions of those born at the extremes of birth weight are likely to differ substantially from each other. Despite this, the DNA methylation profiles of infants at both ends of the growth spectrum are more similar to each other than to the control subjects. Several factors contribute to the strength of this study. First, the two-stage design of this study increases confidence in our findings. In a recent review of 257 epigenome-wide association studies (EWAS) the median number of study subjects included was 46, with only about one third of studies validating results in a second cohort^21^. Our technical verification and then validation in Cohort 2 illustrates the robustness of the predictions from the genome-wide assays, increasing confidence in our results. We tested CD34+ HSPCs as cells with both the long-lived properties and mechanistic properties in inflammation and maintenance of vascular integrity that make it a potential mediator of adult disease risks associated with extreme fetal growth.

We find that IUGR and LGA share a common response with a tendency toward increased DNA methylation. The targeted loci show enrichment at candidate *cis*-regulatory elements and proximity to genes encoding proteins with functions implicated in the *Maturity onset diabetes of the young* and *Hedgehog (HH) signaling* pathways, in both IUGR and LGA subjects despite sharing only a subset of identical loci undergoing DNA methylation dysregulation. These gene/protein properties are significant when considered in terms of the adult phenotype associated with abnormal fetal growth, such as premature glucose intolerance and type 2 diabetes mellitus^36^. Hedgehog signaling is critical for stem cell proliferation and self-renewal^37^, is necessary for hematopoietic stem cell fate decisions^38^ and may play a critical role in CD34+ cells reparative contributions after myocardial infarction^39^.

We also find a sexual dimorphism in the DNA methylation profiles, with IUGR males and LGA females showing greatest alterations in global DNA methylation. Some of the large epidemiological studies that examined outcomes for males and females separately have found sex-specific differences^2,40,41^, though these have not been consistently reported. Results from other EWAS studying epigenetic responses to adverse intrauterine conditions have generally presented combined observations for males and females^18,19^. Our findings of a global shift towards increased DNA methylation is inconsistent with findings seen when the relationship between fetal growth and DNA methylation of Long Interspersed Nuclear Elements (LINE-1) was tested in cord blood leukocytes^15,42^ and placenta^14^. Decreased LINE-1 methylation, which has been associated with genomic instability and cancer risk^43^, was found in cord blood from newborns with low and high birth weight^42^. The development of adiposity in 5-12 year old boys but not girls has also been associated with decreased LINE-1 methylation in peripheral leukocytes^44^. Others have examined sex-specific changes in association with other environmental exposures, but these were either in a limited number of differentially methylated regions^45^ or global changes with limited sample size numbers^46,47^. All of the prior studies included mixed cell-type samples, which can hamper interpretation of DNA methylation studies^20^.

To avoid the possibility of artifactual results stemming from testing of mixed cell populations, we examined purified cell samples. CD34+ HSPCs were chosen for their self-renewal properties that enable them to propagate a cellular memory of temporally remote events, and for their mechanistically plausible contribution to the associated adult phenotype, especially the increased susceptibility to cardiovascular complications. CD34+ HSPCs play a key role in maintaining the intravascular endothelial layer. Intimal denudation generally precedes the development of atherosclerosis^48^ and HSPCs contribute to repair after peripheral ischemic injury through differentiation into endothelial cells^49^ but may also mediate repair through stem cell paracrine effects^50,51^. In adults, circulating numbers of CD34+ HSPCs have been shown to be inversely related to cardiovascular disease risk^23,24^.

Furthermore, HSPCs can be induced to differentiate into multiple tissue types, including those involved in metabolic regulation^52,53^. Impaired mobilization of HSPCs from the bone marrow^54^ and decreased circulating HSPCs^55^ are thought to link metabolic disorders such as diabetes to cardiovascular disease risk. The effect of aging on HSPCs includes reduced self-renewal capacity^56,57^ and increased myeloid-biased differentiation^56,57^, which has been associated with the increased susceptibility to chronic age-related diseases^58,59^. HSPCs from young and old individuals are similarly effective in reconstituting blood lineages after transplantation^57,60^, but aged HSPCs may be less effective at homing and engrafting at the sites of injury^56^. The regenerative potential of hematopoietic stem cells is multifaceted and the contributing roles of functional defects in the stem cell population itself versus impairment of the tissue environment are as yet unknown. Future studies are needed to determine the clinical impact for stem cell transplantation when using cord blood samples from these otherwise healthy neonates with abnormal fetal growth, as our study suggest that HSPCs may decrease their abilities to renew and differentiate after exposure.

Studying the epigenetic basis for developmental origins of adult disease poses several challenges due to the inaccessibility of the human fetus *in utero*, the lack of tools to measure intrauterine exposures over course of the pregnancy and the duration of the time needed to study outcomes that evolve over decades of the human lifespan. While no study is without limitations, we present our work in part as a framework for discussion of the challenges and considerations when designing EWAS in the future. Our findings indicate that sex-specific differences should be examined in addition to a range of clinical phenotypes or experimental intrauterine exposures in animal models. CD34+ HSPCs represent a homogenous, accessible cell population directly relevant to the study of developmental origins of adult disease given their known involvement in cardiovascular disease risk. CD34+ HSPCs are also well characterized from a genomic perspective through Roadmap Epigenomics Program mapping studies, allowing the investigator to interpret findings in terms of functional elements in this cell type specifically.

Defining the functional implications of perturbations in the pathways identified here will be a valuable further avenue of research. Our findings provide key insights into how seemingly opposing intrauterine exposures give rise to similar adult phenotypes, through perturbation of DNA methylation converging on common loci or at distinct loci targeting genes in common pathways. As methods for design and execution of EWAS become better defined, the discovery of novel biomarkers that represent cumulative prior exposures in early life may ultimately provide new tools that identify at-risk neonates for preventative interventions.

## METHODS

This study was approved by the institutional review board (IBR) of the Montefiore Medical Center and the Committee on Clinical Investigation at the Albert Einstein College of Medicine and is in accordance with Health Insurance Portability and Accountability Act (HIPAA) regulations. Written informed consent was obtained from all subjects prior to participation.

### Sample Collection

Cord blood from neonates was the source of material for this study. Biological samples and clinical information were collected (n=84) from consenting women who delivered healthy infants without any anomalies or dysmorphic features and following an uncomplicated intrapartum course, without evidence of fetal distress (normal Apgar scores and cord blood gases without acidemia). The three groups were comprised of infants with appropriate growth, IUGR or LGA (matched for gestational age at delivery and sex). Both birth weight and ponderal index (a measurement of neonatal weight relative to length) were used to identify case and control subjects. IUGR and LGA were respectively defined by birth weight and ponderal index values <10^th^ or >90^th^ percentile for gestational age and sex. Control infants had normal parameters (>10^th^ and <90^th^ percentiles) for both birth weight and ponderal index.

Maternal and infant characteristics are shown in **Table 1**. Cohort 1 (genome-wide assays) is composed of 20 samples per group, while Cohort 2 (validation cohort, targeted assays) has 8 samples per group, with equal representation between male and female subjects in all groups.

### Isolation of CD34+ HSPCs

CD34+ cells, which constitute approximately 1% of nucleated blood cells in umbilical cord blood^61^, were isolated from the cord blood specimen using an immunomagnetic separation technique. Mononuclear cells were separated by Ficoll-Paque density gradient or using PrepaCyte-WBC following which CD34+ cells were obtained by positive immunomagnetic bead selection, using the AutoMACS Separator (Miltenyi Biotech). This resulted in the isolation of cells with ≥95% purity. We cryopreserved the purified cells in 10% DMSO using controlled rate freezing.

### Genome-wide DNA methylation assay

The HELP-tagging assay was performed after isolation of genomic DNA from frozen CD34+ HSPCs, digested to completion by either HpaII or MspI. The digested DNA was ligated to two custom adapters containing Illumina adapter sequences, an EcoP15I recognition site and the T7 promoter sequence. Using EcoP15I, we isolated sequence tags flanking the sites digested by each enzyme, methylation-sensitive HpaII or methylation-insensitive MspI, followed by massively-parallel sequencing of the resulting libraries (Illumina technology)^26^. HpaII profiles were obtained for each sample (n=60), calculating methylation scores using a previously generated MspI human reference.

### Data processing and statistical analysis

DNA methylation scores from 0 (fully methylated) to 100 (unmethylated) were filtered by confidence scores. These confidence scores were calculated for each sample based on the total number of HpaII-generated reads as a function of the total number of MspI-generated reads, excluding loci for which the confidence score was lower than the expected mean by locus. To understand the relative effects of known technical covariates acting on methylation data variability, we performed principal components analysis (PCA, R package *princomp*) on the DNA methylation score obtained from the preprocessed data. We fit a linear model for each of the 10 principal components as a function of each covariate, and summarized the data with a heatmap of the negative log_10_ p-values of each regression. We found batch effects (date of sequencing and presence in the same lane of the Illumina machine) to be significant confounding covariates **(Supplementary Fig. 7)**. We confirmed that the effect of a global increase of DNA methylation in cases compared to controls remained after controlling by adjusting p values for the batch covariate.

Candidate differentially methylated loci were identified using ANOVA with pairwise two-tailed Tukey-tests when comparing controls with either IUGR or LGA as well as two-sided t tests when comparing control/cases to define locus specific differences in average methylation between groups. Confirmatory linear regression of DNA methylation on group adjusting for batch effects was also performed. Comparisons between control/IUGR, control/LGA, and IUGR/LGA were also stratified by sex. For each comparison, only loci with at least 8 samples from each sex were retained. After confidence score filtering and selection for a minimum number of observations in each group, the number of testable loci decreased from >1.8 million to 993,514. We defined candidate differentially methylated loci to have a difference between mean DNA methylation scores >20% and a p value <0.05. The necessary amplitude of DNA methylation score differences was defined using power calculations from a preliminary analysis of a subset of our samples (5 per group). Using the average methylation from the control group, the standard deviations from each of the three groups and 16 samples per group (our minimum sample requirement per group after QC), we are fully powered (>99%) to detect at least one group methylation difference >20% at an FDR=0.05. To demonstrate that our technique still exceeds minimum power recommendations for gender specific comparisons, we ran simulations of the gender comparisons with 8 samples per group and we were powered at roughly 85% to detect at least one group methylation difference >20% at an FDR=0.05.

### Bisulphite MassArray verification assays

We selected 24 samples from Cohort 1 to test the technical performance of the genome-wide DNA methylation studies, a verification approach on our original cohort. Bisulphite conversion and MassArray (Sequenom, San Diego, CA, USA) were performed^63^. Primers were designed to cover loci with low, intermediate and high levels of DNA methylation from the HELP-tagging data across all samples regardless of group (**Supplementary Fig. 4**).

### Targeted bisulphite sequencing (TBS)

We bisulphite-converted 200 ng of DNA using the Zymo EZ-96 Methylation-Lightning Kit. After separate PCR amplification of individual target regions (primers listed in **Supplementary Table 5**), we pooled the amplicons in equal ratios and generated Illumina libraries using robotic automation (Tecan). In total, 24 libraries were multiplexed on the Illumina Miseq for 250 bp paired end sequencing. Amplicons were selected to be part of our differentially methylated loci, covering the entire spectrum of predicted DNA methylation values (from 0 to 100) and allowing us to validate our specific pathways.

### Amplicon bisulphite sequence alignment, DNA methylation calls

Sequence reads from the Illumina MiSeq were trimmed for adapter sequences and aligned to the human genome using *bsmap* (Bisulphite Sequencing Mapping Platform)^64^ using the default settings, requiring a PHRED score of ≥37 during alignment. We checked for bisulphite conversion efficiency (C → T in CH contexts, **Supplementary Table 6**) and quantified the percent methylation for each sample (from 0 (unmethylated) to 1 (fully methylated)) at every CpG in the amplicons using the *methratio* tool provided by *bsmap*. We performed validation on 24 new CD34+ HSPC samples (Cohort 2 with 8 subjects/group).

### Genome annotation

We obtained publicly available chromatin immunoprecipitation followed by massively-parallel sequencing (ChIP-seq) data from the Roadmap in Epigenomics project for CD34+ mobilized HSPCs from a 33 year old, Caucasian female (RO_01549/GSM706857). Annotation of genomic features consisted of processing raw data provided through http://www.roadmapepigenomics.org/ for chromatin accessibility (DNase hypersensitivity) as well as ChIP-seq data for six histone modifications, followed by the use of the Segway algorithm^28^ to predict 7 features, interpreted using Self-Organizing Maps^29^ and RefSeq gene metaplots to define promoter, enhancer, transcribed and repressed sequences in the CD34+ HSPCs.

### Functional enrichment analysis

To perform gene set enrichment analysis (GSEA)^65^ we first linked RefSeq genes to our candidate differentially-methylated loci. We filtered these candidate differentially-methylated loci to include only those overlapping candidate promoters (feature 6) or enhancers (feature 4), thereby enriching for loci with greater likelihood to have functional consequences. Candidate differentially-methylated loci overlapping candidate promoters within ±2 kb and candidate enhancers within ±5 kb of RefSeq gene transcription start sites were used to link DNA methylation changes with specific genes. Differentially enriched pathways found using a False Discovery Rate (FDR) q value <0.05 are shown in **Supplementary Table 1**. We validated the KEGG *Maturity onset diabetes of the young* and *HH signaling* pathways (**Supplementary Table 2**) using the Bioconductor package *GOseq*^32^ to control for bias due to the variation of number of HpaII sites associated with different genes. As our analyses included a large number of genes, we wanted to test further the robustness of the enrichment of the 2 pathways selected, by generating random datasets using a permutation approach. As our original analysis was based on the top 2,000 candidate differentially-methylated loci from HELP-tagging, we selected 2,000 genes randomly from those represented by HELP-tagging (97.6% of total) from the hg19 RefGene database (R package *geneLenDataBase*, database org.Hs.egREFSEQ2EG) 1,000 or 3,000 times, and ran the *GOseq* algorithm on each of these samples. The *Maturity onset diabetes of the young* pathway was not predicted in any of the 1,000 iterations, and 3 times in the 3,000 iterations, while *Hedgehog signaling* was predicted once in the 1,000 iterations and not at all in the 3,000 iterations. We therefore define the observed enrichment for these pathways at our dysregulated genes to be specific and statistically significant (p<0.001). We visualize the association of DNA methylation changes and gene properties using pathways identified from the Kyoto Encyclopedia of Genes and Genomes (KEGG, http://www.genome.jp/kegg/pathway.html). Gene pathways were visualized in Cytoscape v3.0.2 with edges representing the physical interactions between nodes (genes/proteins). Node colors and sizes were adjusted to reflect the enrichment for IUGR or LGA separately or together (Fig. 4), and for sex specificity (**Supplementary Fig. 3**). A complete list of genes associated with these pathways is shown in **Supplementary Table 7**.

## ACCESSION CODES

All HELP-Tagging data have been deposited in the Gene Expression Omnibus (GEO) database under accession code GSE53177.

## ACKNOWLEDGMENTS

Support for this project was provided by the Roadmap Epigenomics Program, R01 HD063791 (Einstein/Greally). Support was also provided by Einstein’s Medical Student Training Program (to NAW and JNT, NIH NIGMS T32 GM007288), the Reproductive Scientist Development Program (to HJH, NICHD K12HD00849-26), and Einstein’s Center for Epigenomics, including the Epigenomics Shared Facility and Computational Epigenomics Group.

## AUTHOR CONTRIBUTION STATEMENT

FD, NAW, JNT, HH: performed experiments, analyzed data. YMZ: performed experiments. NAW, JMG:guided and performed analytical approaches. FD: wrote manuscript. FHE, JMG: designed study, analyzed data, and wrote manuscript.

